# High environmental temperatures put nest excavation by ants on fast forward: they dig the same nests, faster

**DOI:** 10.1101/2025.03.24.645078

**Authors:** Alann Rathery, Giulio Facchini, Lewis G. Halsey, Andrea Perna

**Affiliations:** University of Roehampton, School of Life and Health Sciences, London, UK; IMT School for Advanced Studies, Lucca, Italy; Laboratoire Matière et Systèmes Complexes, CNRS, Université Paris Cité, Paris, France

**Keywords:** Ants, digging, temperature, networks, pattern formation

## Abstract

Environmental temperature influences the physiology and the behaviour of ectothermic organisms, including ants. However, the complex collective behaviour exhibited by ant colonies means that it is difficult to predict how the effects of temperature translate to colony-level functioning and features, such as the form of their nests. This study aims to determine the effects of environmental temperature on nest excavation rate and on the morphology of excavated nests. To this end, we characterized the nest digging activity of the yellow meadow ant *Lasius flavus* confined to dig in a nearly two-dimensional experimental setup maintained at a constant temperature ranging from 15 to 30 degrees Celsius. Ants dug faster at higher temperature, with an increase of digging rate that reflected the temperature-induced increase of movement speed of individual ants. Nevertheless, the shape of excavated nests remained statistically unchanged across the full range of temperatures we tested. These results suggest that temperature accelerates all aspects of the excavation process uniformly, rather than selectively influencing specific components such as tunnel branching or elongation. The ability to produce a consistent overall nest structure, irrespective of the temperature conditions encountered at the time of digging, may provide adaptive benefits to the colony.

## Introduction

Most ant species modify the environment in which they live by building, or digging, structures such as nests, mounds, underground galleries and trails (Hansell 2005; Tschinkel 2021; Perna and Latty 2014; Perna and Theraulaz 2017).

These structures provide protection for the colony, channel the movement of individual ants towards food and other resources, and indirectly determine the activities that they perform, with implications for division of labour among colony members, the organisation of foraging and the spreading of pathogens and diseases within the colony. (Pinter-Wollman 2015; Vaes et al. 2020; Lehue and Detrain 2020; O’Fallon 2023; Pie et al. 2004; Leckie et al. 2024).

Because of their importance for the colony, the shape and size of ant-made structures should be finely controlled, ensuring that their adaptive value is preserved, irrespective of the environmental conditions encountered at the time of construction. However, we know that the size and shape of ant nests is influenced by environmental factors, such as temperature gradients (Sankovitz & Purcell, 2021; García et al., 2024) and humidity (DiRienzo & Dornhaus, 2017). This is likely because environmental factors influence the digging behaviour of ants and therefore, indirectly, the collective output of their behaviour (Detrain & Deneubourg, 2006). For example, the workers of the desert ant *Messor ebenius* tend to dig toward areas of higher water concentration (Tohmé 1972). CO_2_ levels are also known to elicit ant digging activity (Hangartner, 1969), and can contribute to orienting the movement and activity of ants (Römer et al., 2017; Römer et al., 2018).

Ants themselves are ectotherms, meaning that their metabolism and activity are expected to vary with environmental temperature (Gillooly et al., 2001; Dell, 2011; Abram et al., 2017). This means that environmental temperature is also likely to impact ant excavation activity, and in principle could affect the characteristics of excavated nests.

Here, we investigate how digging activity is affected by environmental temperature, through study of the collective digging behaviour of yellow meadow ants, *Lasius flavus*, in laboratory conditions at controlled ambient temperatures (ranging from 15 to 30 °C). We confine experimental populations of our ants to dig in a nearly 2-dimensional layer of clay and sand, which allows us to measure nest growth and characterise nest shape over time. Digging is a complex activity that involves multiple actions (collection, transportation, deposition of pellets) and involves collective-level regulations. To contrast the effects of temperature on digging, against temperature effects on simpler unspecific tasks such as muscular activity and locomotion, we also measure the walking speed of individual *L. flavus* ants across different temperature conditions. We aim to establish if – and how – nest-digging rate and the shape of excavated galleries are affected by environmental temperature.

### Methods Ant colonies

We collected four colonies of *Lasiu*s *flavus* (approx. 2,500-3,000 workers each), from locations in Putney Heath, London, United Kingdom (51° 27’ 0” N; 0° 14’ 11” W). After collection, the colonies were housed in separate plastic trays at controlled room temperature (22 °C) with natural illumination but away from direct sunlight. The ants were provided with dark test tubes half-filled with wet cotton that they could use as shelters, and had *ad libitum* access to water and food. Food consisted of commercial ant jelly food (from *Ants UK*) and honey mixed with water as sources of energy, complemented by sources of protein such as boiled eggs, mealworms, and tuna. All colonies comprised workers and brood, but no queens. At the beginning of each experiment, 200 workers were collected from one of the four colonies and introduced into the digging setup.

### Digging setup

The digging setup consisted of a 2-mm layer of substrate (a mix of 75 % sand and 25 % clay moistened with tap water at approximately 15 % of the dry weight, following Toffin, 2009). The substrate was held horizontally between two A4-sized sheets of glass, stuck together with a rubber band to avoid the desiccation of the substrate. On top of the upper sheet, a plastic box housed the experimental population of 200 ants, and was connected to the substrate through a small hole in the glass and a plastic tunnel of radius 4.5 mm. Multiple digging setups of this type were produced. After introducing 200 ants, each setup was placed into a thermal cabinet at a constant temperature of either 15, 20, 25 or 30° C. The digging setups were continuously exposed to constant artificial light. A camera placed approximately 25 cm below the setup, and controlled through a Raspberry Pi single board computer, recorded images of the setup every 10 min. Each experiment ran for four days at which point the setup was removed from the thermal cabinet. We replicated this experiment multiple times for each colony and at each experimental temperature, for a total of 58 experiments (i.e. an average of 3.6 replicas per colony and temperature condition). Ant mortality was very low across all experiments (< 1 %), with no detectable effect of temperature.

### Image analysis

All recorded images were automatically processed with a custom macro using the software *ImageJ*. Each image was converted to grayscale and excavated galleries were segmented by applying a threshold on the intensity level of the pixels. The segmented images were further cleaned by applying a morphological ‘opening’ operation to remove small, isolated pellets. For each image, the total excavated area was measured by counting the number of pixels identified as galleries, and converting from units of pixels to cm^2^.

For each experiment, the increase of excavated area over time was approximately linear, except for an initial lag phase immediately after ants were introduced in the setup (before ants entered the digging area in-between the two layers of glass), and for a reduction of digging activity near the end of the experiment (possibly due to fatigue, or to the layer of sand and clay starting to lose humidity). Digging rate was estimated by fitting a straight line through the part of the curve with an approximately linear increase of excavated area, and determining the slope of the regression in units of cm^2^/h.

### Nest shape analysis

We characterised nest shape in terms of (a) the shape of the individual tunnels and (b) the larger scale organisation of the network of galleries. We refer to a tunnel as the unbranched portion of a gallery in-between two junctions, or between a junction and an endpoint. Tunnels were described by their *length*, their *width*, and their *aspect ratio* (the ratio of length over width). A network representation of the excavated structures was obtained by mapping junctions and endpoints of each tunnel onto nodes, and the tunnels themselves onto edges. The shape of each network was then summarised in terms of two measures that describe the density and redundancy of connections, *network density* and *meshedness*, and two measures that quantify the length of distances to be travelled to traverse the network, *network efficiency* and *diameter* (see table 1).

**Table 1.** Network descriptors used in this study. The distances d_ij_ are metric distances (cumulative lengths of all edges along the path). In these formulae, m represents the total number of edges in the network and n is the number of nodes

| Measure | Formula | Description |
| --- | --- | --- |
| Diameter | $\max(d_{ij})$ | The diameter is the network-based distance between the two nodes that are farthest apart in the network. It describes the spread of the network. $d_{ij}$ is the shortest network-based distance between node $i$ and node $j$. |
| Density | $\frac{m}{(n(n-1))/2}$ | The number $m$ of observed edges in the network relative to the number of edges in the corresponding fully-connected network (the network with the same number nodes $n$, and where there is a direct connection between all pairs of nodes). |
| Network Efficiency | $\frac{1}{n(n-1)} \sum \frac{1}{d_{ij}}$ | Efficient networks have short paths between pairs of nodes. The measure of network efficiency averages the inverse of the shortest network-based distance ($d_{ij}$) over all pairs of nodes. |
| Meshedness | $\frac{m - n + 1}{2n - 5}$ | Number of observed cycles in the network, relative to the maximum number of cycles in a planar graph with the same number of nodes. |

The mapping of the network was done by skeletonizing the segmented images in *ImageJ* (*Skeletonize* plugin). The width of each tunnel was calculated as the average width over all the pixels on the image skeleton of the tunnel, calculated via a *Chamfer* distance algorithm in *ImageJ*.

### Ant speed

In a separate series of experiments, we measured the movement speed of individual ants. A sample of worker ants from the same colonies used for the digging experiments were tested individually in empty Petri dishes, the vertical edges of which had been coated with Fluon. The Petri dish, with the ant, was placed in the thermal cabinet at constant temperature within the range 12.5 to 32.5 °C. The Petri dish was recorded at 10 frames/s with a camera controlled by a Raspberry Pi computer. Each recording lasted for one hour. The ant position was automatically tracked throughout the video with a custom-made script written in *Python* (which called the MOG background subtraction method of the *OpenCV* library) and the speed of the ant was calculated. Periods that the ant spent motionless or slowly moving were automatically excluded from the tracking for technical reasons (the tracking only detects objects that move in the image) and for biological reasons (the proportion of time that an ant spent motionless can be variable). Each ant was tested at a single temperature to avoid possible cumulative effects from repeated experiments. 218 ants in total were tested in this way.

### Data analysis

The effect of environmental temperature on digging rate was quantified by fitting the Sharpe-Schoolfield equation implemented in the *rTPC* package in *R* (Sharpe and DeMichele, 1977; Schoolfield *et al*., 1981; Padfield *et al*., 2021) to the measured digging rates of each colony. The same equation was also fitted to the data of individual ant movement, to quantify the effect of temperature on walking speed.

The Sharpe-Schoolfield equation assumes that biological rates *R*(*T*) increase with temperature following an Arrhenius-type relation, and then rapidly decrease at very high values of temperature. In the results that we report here, we focus on the Arrhenius part of the curve, which is well approximated as follows:

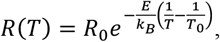

and is described by two parameters: a value of activation energy *E* (measured in electronvolts, eV), which describes how fast the rate increases with increasing temperature, and a reference rate *R*_*0*_, which is the rate of the process at some (arbitrarily chosen) reference temperature *T*_*0*_ (throughout all our analyses we set *T*_*0*_ = 293.15 K; *k*_*B*_ is the Boltzmann constant).

Additional terms of the Schoolfield equation not reported here (but detailed in the cited references) allow the Sharpe-Schoolfield equation to also fit the plateauing and then the decrease of biological rates at high temperatures. In our study we did not focus on testing individual and collective ant behaviour at temperatures higher than 30 °C, when ants could be stressed or die. To force the Sharpe-Schoolfield model to fit the data despite the lack of high-temperature measurements, we added one extra data point to all datasets, corresponding to the unmeasured assumption that ant digging and walking activity stop at 43 °C (which is the maximum critical temperature CT_max_ of *Lasius flavus*, Corley et al., 2023).

We analysed the shape of nests at two values of size for each nest: when the excavated area reached 50 cm^2^ and then when it reached 100 cm^2^. We made the choice to analyse shapes at a fixed size (rather than a fixed time of excavation) because the comparison of network descriptors is only meaningful for similarly sized networks. The specific size values of 50 cm^2^ and 100 cm^2^ were chosen because they are sufficiently large that structures offer several measurable features, but small enough that excavated galleries have not yet reached the physical boundaries of the digging area (where interactions with the boundary could alter the shape). Nevertheless, a small proportion of experiments (typically those with a long initial lag-phase, in which ants had remained aggregated outside for too long before entering the digging area) failed to reach the set sizes and by the end of the experiment and could not be measured: we measured nest features on 51 nests that had reached the excavated area of 50 cm^2^ and 39 of the same nests that had also reached the excavated area of 100 cm^2^.

For the comparison of tunnel shape and network descriptors across different temperature conditions we ran linear mixed-effects models using the *lmer* function (*lme4* package) in *R* (Bates et al., 2015). We considered temperature as a continuous variable and the excavated area as a fixed factor, with the ant colony excavating the nest included as a random factor and the individual ID of the experiment replicate (corresponding to the fact that nests were measured at 50 and 100 cm^2^) nested into colony:

`experimental_variable ~ temperature + area + (1|colony/ID)`.

Throughout the text, we report the results of fitting these models in terms of fixed effects coefficient β_T_ (for temperature) and β_A_ (for nest area), and in terms of the associated standard error SE, and P-value. We fitted models to all the different nest descriptors indicated in the *nest shape analysis* section above. We did not apply corrections for multiple statistical tests. This is because correcting for multiple tests reduces the chances of a type I error, but necessarily increases the chances of a type II error, i.e. reporting no significant difference when a significant effect is in fact present. We preferred, therefore, to not correct for multiple testing and instead, we interpret p-values arising from testing with caution.

All analyses were performed in *R*, version 4.3.2. To minimize experimental bias, tests at different temperatures were carried in randomized order, and data analysis relied on the same automated scripts for all tested conditions.

## Results

Ants consistently entered the digging substrate and excavated complex patterns of branching and reconnecting galleries (fig. 1).

**Fig 1.**
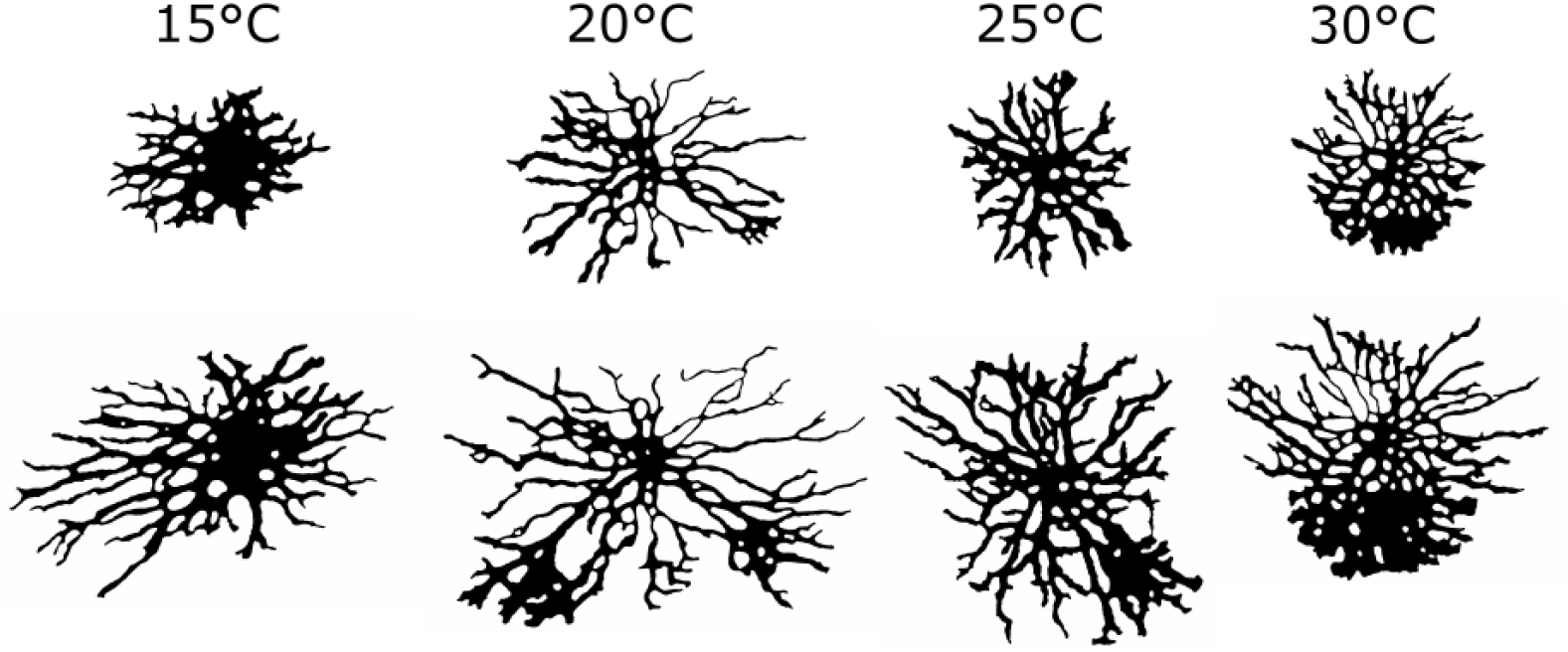
Examples of segmented galleries from four experiments at different temperatures. The top and bottom image in each column represent the same excavated structure at two different sizes (top: 50 cm^2^; bottom: 100 cm^2^)

The digging rate increased with environmental temperature for all colonies and throughout the range of temperature conditions that we tested (15 to 30 °C, fig. 2), plateauing around 30 °C: the best-fitting Sharpe-Schoolfield equations reached the maximum digging rate between 25 and 30 °C in two of the four colonies, while for the other two colonies, the fit was still increasing at 30 °C. The fitted values of activation energy ranged between 0.23 and 0.88 eV across colonies (when data from all colonies were analysed together, they led to a value of activation energy of 0.49 eV; 95 % CI [0.33, 1.60]). A linear model of digging rate as a function of temperature in Celsius degrees, with colony identity as a random factor indicates a strong effect of temperature on digging rate (β_T_=0.182, SE=0.030, P<0.01).

**Fig 2.**
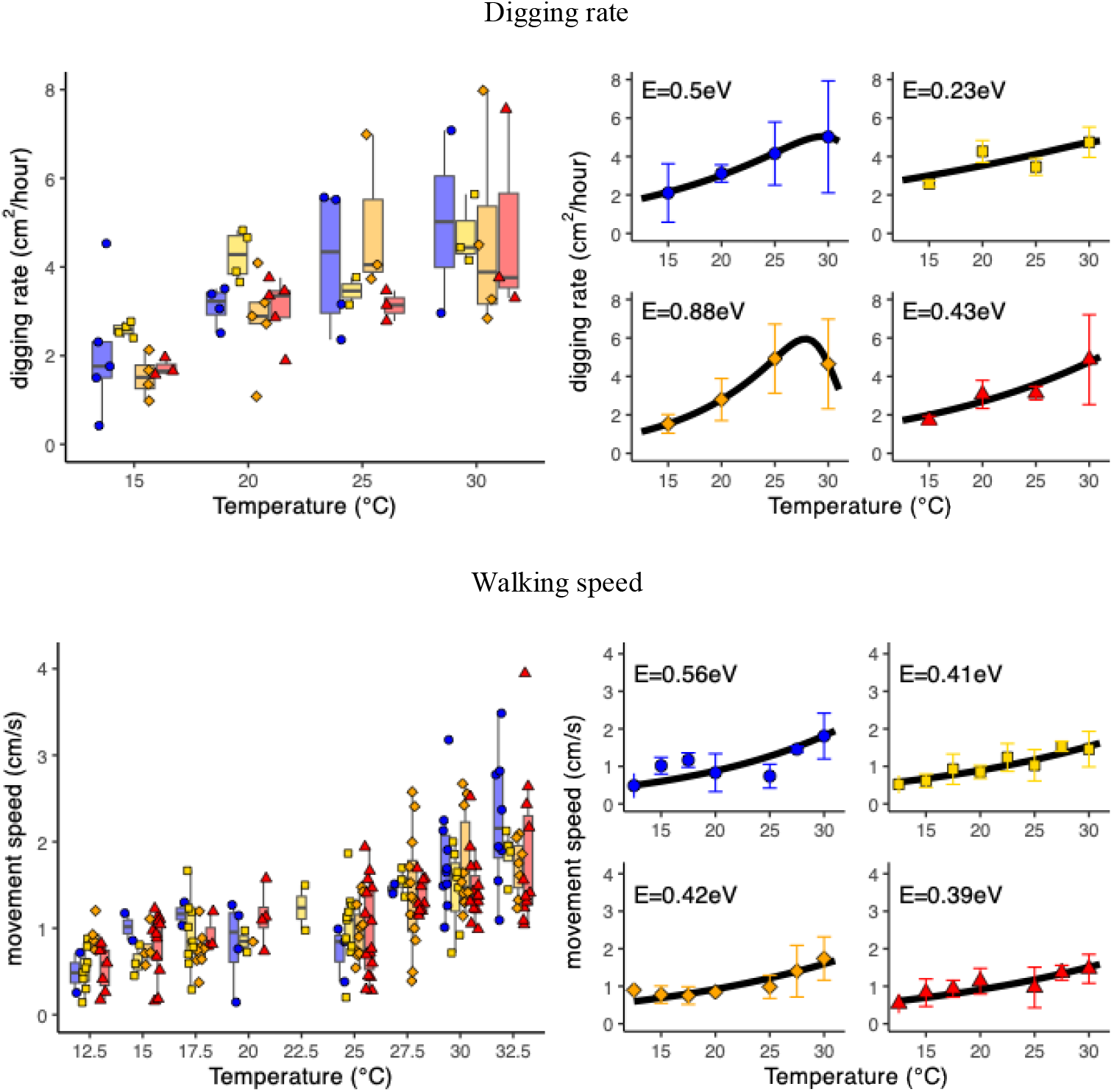
Top panels. Effect of temperature on ant digging rate. Left: measured digging rates, as a function of environmental temperature, for each colony and replica of the experiment. Each marker corresponds to one replicate of the experiment; marker colour and shape indicate the identity of the original ant colony. Right panels: fitted Sharpe-Schoolfield equations (in black) for each colony; markers and error bars represent the mean digging rate ± 1 standard deviation across replicates for one colony. The text on each graph reports the fitted value of activation energy. Bottom panels. Effect of temperature on the walking speed of individual ants. Left: each marker corresponds to the movement speed of one individually tested ant (75th percentile of the speed distribution for that particular ant); marker colour and shape indicate the identity of the colony of origin of the ant. Right panels: fitted Sharpe-Schoolfield equations for each colony; markers and error bars represent the mean walking speed ± 1 standard deviation across replicates for one colony

**Fig 3.**
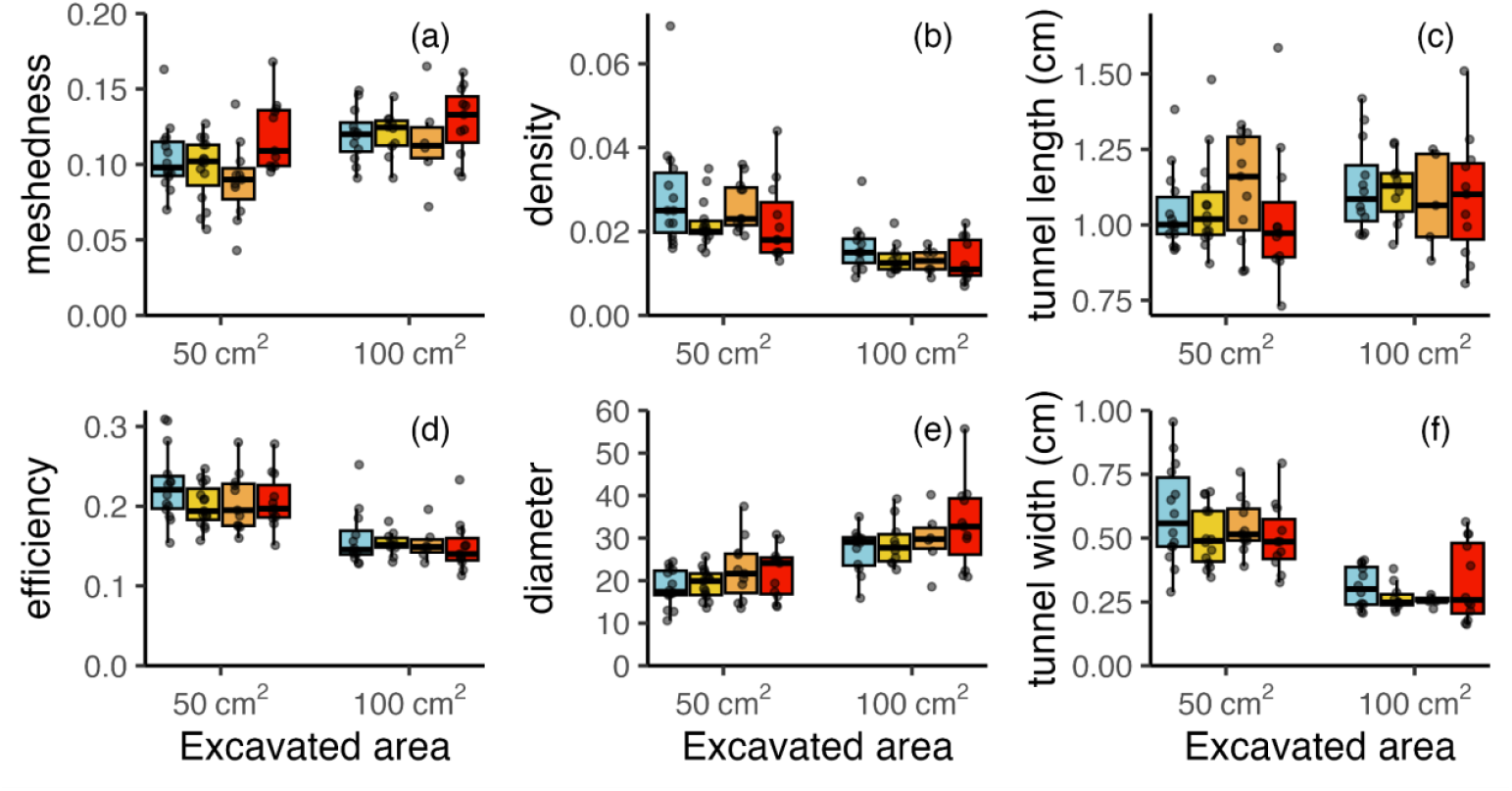
Effect of temperature and excavated area on nest features. (a) network meshedness, (b) network density, (c) tunnel length, (d) network efficiency, (e) network diameter, (f) tunnel width. Each individual data point corresponds to the measurement from a single experiment and at one value of nest size (50 cm^2^ and 100 cm^2^). In case of local network properties which are calculated at the tunnel level, such as tunnel width, a data point represents the average value over the entire network). Box colours indicate the temperature condition of the experiment (cyan=15 °C, yellow=20 °C, orange=25 °C, red=30 °C)

For comparison, we also measured the effect of temperature on the walking speed of individual ants. Ants walked faster, on average, at higher temperatures (fig. 2), with values of activation energy in the range 0.39 to 0.56 eV, so similar to that for digging (when data from all colonies were analysed together, they led to a value of activation energy of 0.43 eV; 95 % CI [0.36, 0.52]).

In spite of differences in digging rate, nest morphology remained similar across experimental conditions. The structures excavated at different temperatures were not distinguishable from one another through visual inspection when comparison involved structures matched for size (fig 1). Quantitative analyses confirmed this visual impression – we found no statistically significant effect of temperature on the length of tunnels (β_T_=0.007, SE=0.007, P=0.29). Similarly, there was no statistically significant effect of temperature on tunnel width (β_T_=−0.001, SE=0.002, P=0.76) or aspect ratio (β_T_=0.016, SE=0.029, P=0.58).

Temperature did not have a statistically significant impact on network meshedness (β_T_=0.001, SE=0.000, P=0.21), network efficiency (β_T_=−0.001, SE=0.001, P=0.40) and density (β_T_=−0.000, SE=0.000, P=0.28). The relation of network diameter with temperature was described by β_T_=0.298, SE=0.133, P=0.03.

While temperature did not, in general, affect nest morphology, the shape of excavated galleries changed over time, resulting in differences between the structures with an excavated size of 50 cm^2^ and those measured at 100 cm^2^ (tunnel width: β_A_=0.254, SE=0.018, P<0.001, tunnel length: β_A_=−0.128, SE=0.075, P=0.097, tunnel aspect ratio: β_A_=−3.049, SE=0.218, P<0.001) and network characteristics (meshedness: β_A_=0.015, SE=0.003, P<0.001, efficiency: β_A_=−0.062, SE=0.003, P<0.001, density: β_A_=−0.011, SE=0.001, P<0.001, diameter: β_A_=10.316, SE=0.754, P<0.001; positive values of β_A_ indicate an increase in the quantity between 50 cm2 and 100 cm^2^).

## Discussion

Temperature had a strong effect on ant digging activity, summarised by the values of activation energy in the range 0.23 to 0.88 eV. These values of activation energy can also be summarised by the statement that the ants excavated between 1.4 and 3.2 times faster at 25 °C than at 15 °C (these increases of activity over a 10 degrees interval are often reported under the name of Q_10_ factors and so, in the case of *L. flavus* digging, the Q_10_ is in the range 1.4 – 3.2). Despite this increase in activity with temperature, the shape of the excavated galleries was not detectably different across the entire temperature range.

The fact that temperature affects the digging rate of ants is not surprising, as many aspects of the physiology and behaviour of ants are greatly affected by temperature (Hurlbert et al., 2008; Andrew et al., 2013; Van Oudenhove et al., 2011; Van Oudenhove et al., 2012; Traniello et al. 1984; Tizón et al., 2014). Previous studies have already reported on the relation between environmental temperature and digging activity in various ant species (Bollazzi, Kronenbitter and Roces, 2008; Sankovitz & Purcell, 2021; García Ibarra et al., 2024), typically finding an increase of digging activity with increasing temperature. In these previous studies, variations of temperature were often associated with the presence of spatial temperature gradients - areas of a nest farther from the surface of the ground are less affected by temperature variations. Our study confirms that similar findings also hold when temperature is modulated in the absence of spatial gradients.

We did not detect consistent effects of temperature on nest shape, either at the local scale of individual corridors or at the broader scale of the excavated network of galleries as a whole. Ants excavated similar structures across the entire range of temperatures. A recent study focusing on the effect of temperature and humidity on nest excavation by harvester ants (O’Fallon *et al*., 2022) similarly found that neither temperature nor humidity were inducing changes in the shape of the nests in the absence of gradients in the soil. In the collective behaviour of social insects, ‘shape transitions’ between qualitatively different patterns often occur when two variables that control the process change disproportionately relative to each other. For instance, Toffin and collaborators (2009) characterised a shape transition during nest digging in ants, whereby an initial circular cavity evolves into a ramified structure through a branching process. In their study, the shape transition occurred because the number of workers involved in the excavation remained approximately constant over time while the area available for excavation increased, leading to a decreasing density of ants per available digging site, resulting in a change in the nest shape being created. In the context of responses to temperature, we could expect to observe temperature driven shape transitions if different components of ant digging behaviour had different activation energies, e.g. if the spontaneous initiation of new tunnels was more affected by temperature than the elongation of already existing ones, or if the proportion of digging ants vs. inactive ants changed with temperature. The fact that we did not observe temperature-induced shape differences in our experiments suggests temperature sped up all activities that contribute to the digging process in a similar way, an analogy being when playing a video at normal speed and then sped up. The fact that similar values of activation energy described both ant movement and nest growth supports this conclusion. In conclusion, high temperatures within the normal ecological range for *L. flavus* do not affect how they dig nests, but do result in these ants digging them more quickly. In nature, *L. flavus* ants are typically found in open meadows in temperate regions, and they experience daily and seasonal temperature fluctuations, including inside their mound (see e.g. Véle and Holuša, 2017). There might be an adaptive benefit for the ants in making coherent structures, irrespective of daily and seasonal temperature fluctuations that might be experienced at the time of construction, especially if we consider that some of these structures are likely to persist for considerable time after their initial construction (see e.g. King, 2020).

We should note, however, that while temperature *per se* did not affect nest shape in our experiments, under natural conditions temperature variation is often associated with the presence of gradients of temperature itself and of humidity, and these gradients are known to have an impact on the shape of excavated nests (Bollazzi, Kronenbitter and Roces, 2008; Sankovitz & Purcell, 2021; García Ibarra et al., 2024). Future research could complement our study with a characterisation of galleries excavated by *L. flavus* in their natural field conditions.

In our experiment, we did not reach the thermal limit of *L. flavus* (the thermal limits of the *Lasius* genus are reported to be around 38.7 °C; Diamond *et al*., 2012; Penick et al., 2017). Future work could explore the effect of temperature on digging activity along a wider range of temperature conditions. Our study, which focused on laboratory experiments running for a short period of time, highlights the effects of temperature on digging activity in isolation from other ecological variables such as colony growth and ecological interactions, which are also affected by temperature. For example, a higher temperature is expected to increase the metabolic rate of individual ants, and consequently reduce the population density of ants that can be sustained per unit surface area of land.

In the future, it will be interesting to integrate our results with more empirical information on the effects of temperature on other variables of the behaviour and ecology of *L. flavus* ants. In this way, it might be possible to understand, directly from bottom-up principles, how this ant species, which is an important ecosystem engineer, and the meadows that are so characteristically associated with it, might respond to current and future changes in its environment.

## Data availability

Data and analysis code associated with this manuscript are available at https://doi.org/10.6084/m9.figshare.28602074.

## Authors’ Contribution

AR, GF, LGH and AP conceived and planned the experiments; AR carried out the experiments; GF, LGH and AP provided technical assistance for setting up the experiments and data acquisition; AR and AP analysed the data with input from GF and LGH; AR wrote the initial draft; AP wrote the second draft; GF and LGH provided critical feed-back on all drafts.

## Ethics declaration

Ant collection from Putney Heath was discussed in advance with the local rangers to minimize the impact on human visitors and wildlife. The experimental procedures involved in our study did not require specific ethical approval.

